# The effect of problem-based learning on improving problem-solving, self-directed learning, and critical thinking ability for the pharmacy students

**DOI:** 10.1101/2023.10.26.564146

**Authors:** Yi-Jing Zhao, Feng-Qing Huang, Qun Liu, Ying Li, Raphael N Alolga, Lei Zhang, Gaoxiang Ma

**Author notes:** **Correspondence to:** Gaoxiang Ma, School of Traditional Chinese Pharmacy, China Pharmaceutical University, Nanjing, China.; Lei Zhang, School of Traditional Chinese Pharmacy, China Pharmaceutical University, Nanjing, China. These authors contributed equally to this work (Yi-Jing Zhao, Feng-Qing Huang).

## Abstract

**Objective:** This study aimed to comprehensively evaluate the effect of PBL on problem-solving, self-directed learning, and critical thinking ability of pharmaceutical students through a randomized controlled trial (RCT) and meta-analysis of RCTs.

**Methods:** In 2021, 57 third-year pharmacy students from China Pharmaceutical University were randomly divided into a PBL group and a lecture-based learning (LBL) group. Mean scores were compared between the two groups for problem-solving, self-directed learning, communication skills, critical thinking, and final exam grades. Students’ feedback on the implementation of PBL was also collected. A meta-analysis was subsequently performed on eight studies involving 1,819 students.

**Results:** The PBL group had significantly higher mean scores for problem-solving (8.43±1.56) and self-directed learning (7.39±1.19) than the LBL group (7.02±1.72 and 6.41±1.28, respectively). The PBL group also showed better communication skills (8.86±1.47) than the LBL group (7.68±1.89). The mean level of critical thinking was significantly higher in the PBL group than the LBL group (p=0.02). The PBL group also had better final exam grades (79.86±1.38) compared to the LBL group (68.1±1.76). Student feedback on PBL implementation was positive. The subsequent meta-analysis confirmed these findings.

**Conclusion:** This study found that PBL is an effective teaching method for pharmacy students.

## INTRODUCTION

Pharmacists have a unique position in providing healthcare provision and promotion, which has been demonstrated to lead to favorable health outcomes and decreased overall healthcare costs^1^. To be competent in personalized, flexible, and situation-specific problem-solving, pharmacists require critical thinking and self-directed learning abilities. Critical thinking is a process that involves clarifying, simplifying, organizing, and rationalizing ideas^2^.

Problem-based learning (PBL) has been identified as a useful method for connecting theoretical learning with real-world clinical problems, requiring problem-solving skills, critical thinking, effective communication, utilization of literature, and strong teamwork^3^. PBL is a student-centered learning strategy that uses a problem as a starting point to facilitate learning. PBL allows pharmacy students to collaborate in small groups with the goal of improving their cognitive capacities and clinical and critical thinking skills. PBL has been applied in many healthcare professional programs, including pharmacy, and has been found to improve students’ learning and retention of material beyond the end of the course^4,5^. Although pharmacists have been shown to contribute their clinical expertise and services to healthcare teams^6-8^, challenges still exist regarding the acceptance of this expanded role by other medical professionals^9^. The primary goal of pharmaceutical care is to transition pharmacists from solely dispensing the correct drug to becoming more involved in the selection of optimal drug therapy that is safe and efficacious^9^. In the United States, PBL in pharmacy education has been shown to positively impact students’ abilities to problem-solve and apply a basic science foundation to their clinical decision-making for healthcare treatment^10^.

Several studies have investigated the effects of PBL on academic achievement in pharmacy courses, including final examination and post-test scores, students’ grades, and GPA. Previous meta-analyses have also assessed the impact of PBL on learning achievements in pharmacy students^11,12^. However, these studies have not evaluated the effect of PBL on critical thinking, problem-solving, and self-directed learning ability. Moreover, many of these studies were not randomized, which may introduce bias. To comprehensively evaluate the impact of PBL on critical thinking, problem-solving, and self-directed learning ability in pharmaceutical students, this study performs a randomized controlled trial. Additionally, a meta-analysis of randomized clinical trials will be conducted to assess the outcomes of PBL.

## METHODS

### Study subjects

In the present study, a total of 57 third-year pharmacy students from China Pharmaceutical University were randomly assigned to either a PBL group or a lecture-based learning (LBL) group in 2021. All participants had no prior experience with PBL. Informed consent was obtained from the participants after providing them with a detailed explanation of the study. The data collection instrument was a questionnaire divided into three sections: (1) demographic characteristics, (2) a standard questionnaire assessing student behavior, including problem-solving, self-directed learning ability, communication skills, and critical thinking, and (3) a standard questionnaire assessing student attitudes towards PBL, including students’ role, lecturer role, and the effectiveness of the unit from the students’ viewpoint. The fourth section of the questionnaire included researcher-developed questions aimed at investigating students’ feedback on the implementation of PBL. The protocol was approved by the Committee on Research Ethics of China Pharmaceutical University. The approval number was 2021XJQN14.

### Search strategy and selection for meta-analysis

A total of two databases, PubMed and CNKI, were searched for related studies from inception until December 31, 2022. The search keywords used were “pharmacy OR pharmacology” AND “PBL OR problem-based learning”. The systematic search and data extraction were conducted according to the Preferred Reporting Items for Systematic Reviews and Meta-Analyses (PRISMA) guidelines (**Supplementary Table 1**).

**Table 1.** Demographics of third-year pharmacy students in a study comparing problem-based learning (PBL) and lecture-based learning (LBL) in Authentication of Chinese Medicines.

| Student Demographic | PBL group (n=28) | LBL group (n=29) | *P* |
| --- | --- | --- | --- |
| Gender (%) |  |  | 0.85 |
| Male | 9 | 10 |  |
| Female | 19 | 19 |  |
| Age (mean ± SD) | 21.21±1.24 | 21.17±1.36 | 0.91 |
| Satisfaction with this course (%) |  |  | 0.0005 |
| High | 22 | 8 |  |
| Middle | 5 | 18 |  |
| Low | 1 | 3 |  |
| Interest with this course (%) |  |  | 0.001 |
| High | 23 | 10 |  |
| Middle | 3 | 15 |  |
| Low | 2 | 4 |  |

To minimize differences between studies, we imposed the following methodological restrictions for the inclusion criteria: (1) studies must provide the minimum necessary information about the number of students and related mean and standard deviation of behavior scores; (2) to avoid selection bias, the sample size of each study should be more than 10; (3) the median study time for PBL should be more than one month. If more than one article was published using the same population, we selected the most recent or informative report. Two researchers (Yi-Jing Zhao and Qun Liu) independently assessed the retrieved studies according to the pre-specified criteria, and any discrepancies were resolved by another investigator.

### Data extraction and quality assessment for meta-analysis

The following information was collected for each included study: first author, publication year, country, number of students, mean and standard deviation of behavior score, and study type. Data extraction was performed independently by two authors (Yi-Jing Zhao and Qun Liu) and cross-checked for accuracy.

### Statistical analysis

Descriptive statistics for continuous variables were reported as mean and standard deviation, while frequencies and percentages were used to describe categorical variables. The effect of PBL on critical thinking, problem-solving, and self-directed learning ability of pharmaceutical students was evaluated using a random-effects model in the meta-analysis. Heterogeneity among studies was assessed using the χ^2^ test and I2 statistic. Sensitivity analysis was performed by sequentially removing each study to assess the stability of the results. Begg’s funnel plot and Egger’s test were used to evaluate the presence of publication bias. Statistical significance was set at a two-sided p-value of less than 0.05. All analyses were conducted using the ‘metafor’ package in R software (version R-3.5.1).

## RESULTS

### Evaluating the effect of PBL in ourself cohort

In this randomized controlled trial, a total of 57 students were recruited and randomly assigned to either the PBL group (n=28) or the LBL group (n=29). The groups were comparable in terms of age and gender distribution. Results showed that the PBL group had significantly higher satisfaction and interest levels in the course compared to the LBL group (both *p* < 0.001, **Table 1**).

As presented in **Table 2**, the PBL group had significantly higher mean problem-solving scores (8.43 ± 1.56) compared to the LBL group (7.02 ± 1.72) (*p* = 0.002). The PBL group also had significantly higher self-directed learning scores (7.39 ± 1.19) compared to the LBL group (6.41 ± 1.28) (*p* = 0.004). Additionally, the PBL group had significantly better communication skills (8.86 ± 1.47) compared to the LBL group (7.68 ± 1.89) (*p* = 0.01). The mean level of critical thinking was also significantly higher in the PBL group compared to the LBL group (*p* = 0.02). Furthermore, the PBL group obtained a significantly better final exam grade (79.86 ± 1.38) compared to the LBL group (68.1 ± 1.76).

**Table 2.** The mean score of behaviors for the comparing problem-based learning (PBL) and lecture-based learning (LBL).

| Behaviors | PBL (n=28) |  | LBL (n=29) |  | *P* |
| --- | --- | --- | --- | --- | --- |
|  | Mean | Sd | Mean | Sd |  |
| Problem-solving | 8.43 | 1.56 | 7.02 | 1.72 | 0.002 |
| Self-directed learning | 7.39 | 1.19 | 6.41 | 1.28 | 0.004 |
| Communication skills | 8.86 | 1.47 | 7.68 | 1.89 | 0.01 |
| Critical thinking | 7.96 | 1.55 | 6.84 | 1.96 | 0.02 |
| Final exam score | 79.86 | 1.38 | 68.1 | 1.76 | < 0.001 |

After participating in the PBL, students in the PBL group were surveyed using a 9-question survey with five-point Likert ratings (ranging from 1 = strongly disagree to 5 = strongly agree) to provide feedback. The survey had a participation rate of 100% and was voluntary and confidential. Results from the survey indicated that students perceived the value of PBL for their current or future practices (3.82 ± 0.12) and in collaborating with a team to solve problems (3.93 ± 0.17). Additionally, students perceived that PBL increased their knowledge of how to utilize practice guidelines to support evidence-based recommendations (4.18 ± 1.15) and how to collaborate with a team of pharmacy students to solve problems (3.00 ± 0.26). The aggregate data from the survey are presented in **Table 3**.

**Table 3.**
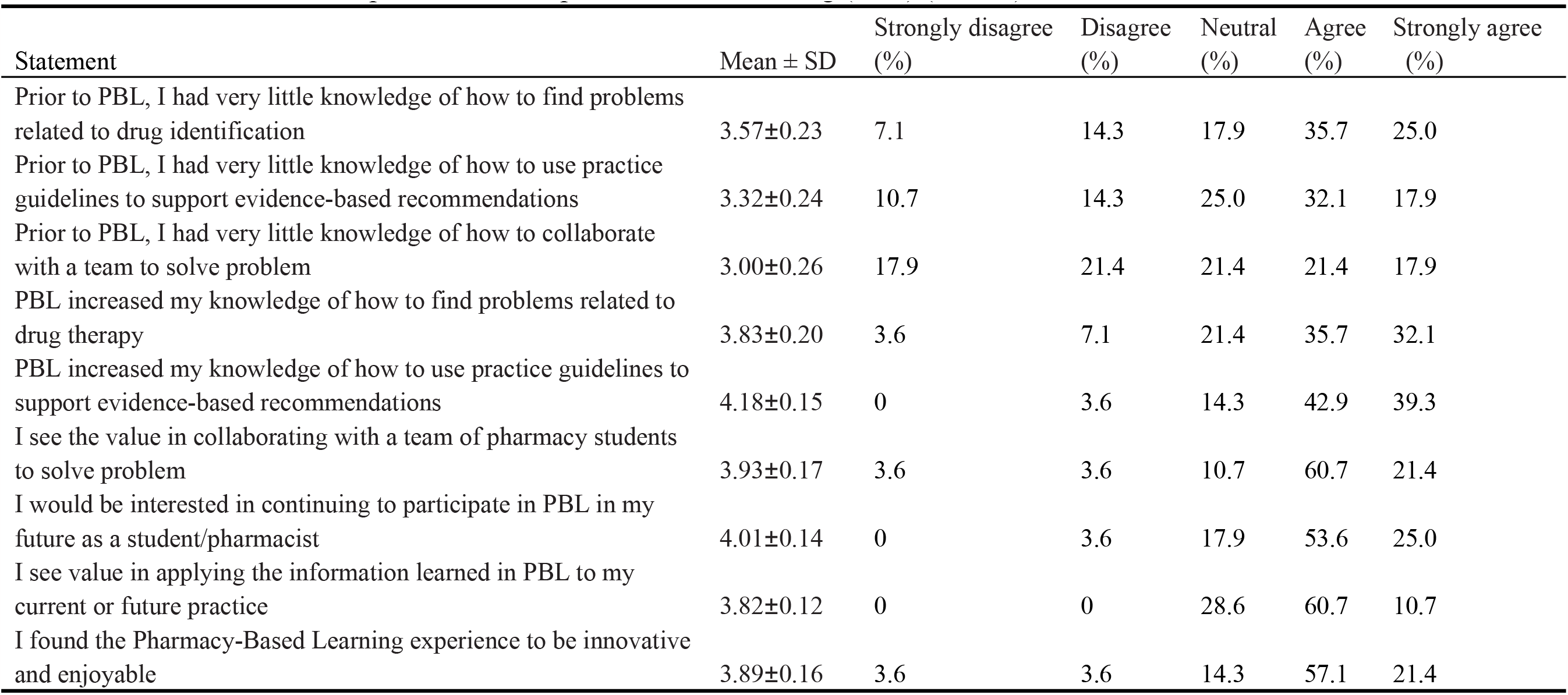
Students feedback on the implementation of problem-based learning (PBL) (N = 28)

| Statement | Mean ± SD | Strongly disagree (%) | Disagree (%) | Neutral (%) | Agree (%) | Strongly agree (%) |
| --- | --- | --- | --- | --- | --- | --- |
| Prior to PBL, I had very little knowledge of how to find problems related to drug identification | 3.57±0.23 | 7.1 | 14.3 | 17.9 | 35.7 | 25.0 |
| Prior to PBL, I had very little knowledge of how to use practice guidelines to support evidence-based recommendations | 3.32±0.24 | 10.7 | 14.3 | 25.0 | 32.1 | 17.9 |
| Prior to PBL, I had very little knowledge of how to collaborate with a team to solve problem | 3.00±0.26 | 17.9 | 21.4 | 21.4 | 21.4 | 17.9 |
| PBL increased my knowledge of how to find problems related to drug therapy | 3.83±0.20 | 3.6 | 7.1 | 21.4 | 35.7 | 32.1 |
| PBL increased my knowledge of how to use practice guidelines to support evidence-based recommendations | 4.18±0.15 | 0 | 3.6 | 14.3 | 42.9 | 39.3 |
| I see the value in collaborating with a team of pharmacy students to solve problem | 3.93±0.17 | 3.6 | 3.6 | 10.7 | 60.7 | 21.4 |
| I would be interested in continuing to participate in PBL in my future as a student/pharmacist | 4.01±0.14 | 0 | 3.6 | 17.9 | 53.6 | 25.0 |
| I see value in applying the information learned in PBL to my current or future practice | 3.82±0.12 | 0 | 0 | 28.6 | 60.7 | 10.7 |
| I found the Pharmacy-Based Learning experience to be innovative and enjoyable | 3.89±0.16 | 3.6 | 3.6 | 14.3 | 57.1 | 21.4 |

### Meta-analysis for the effect of PBL

A total of 4410 articles were searched in the database search. After removing duplicates, 4388 targets were considered for further screening. Following the abstract review, 4039 articles were excluded, and 349 remaining studies were subjected to full-text assessment. After this assessment, 341 studies were excluded based on the following reasons: no available data (n = 305), non-randomized controlled trial research (n = 14), review (n = 8), small sample size (n= 6), and letter and comments (n = 3). Finally, 8 randomized controlled trial articles were included in this meta-analysis (**Figure 1**)^13-20^. As presented in **Supplementary Table 2**, the 8 studies included in this meta-analysis were published between 2002 and 2021 and involved 1819 students from 2 countries. The sample sizes of the included articles ranged from 12 to 1320, and the intervention durations varied from 0.6 to 2 years. The quality assessment of the articles indicated a low or medium risk of bias.

**Figure 1.**
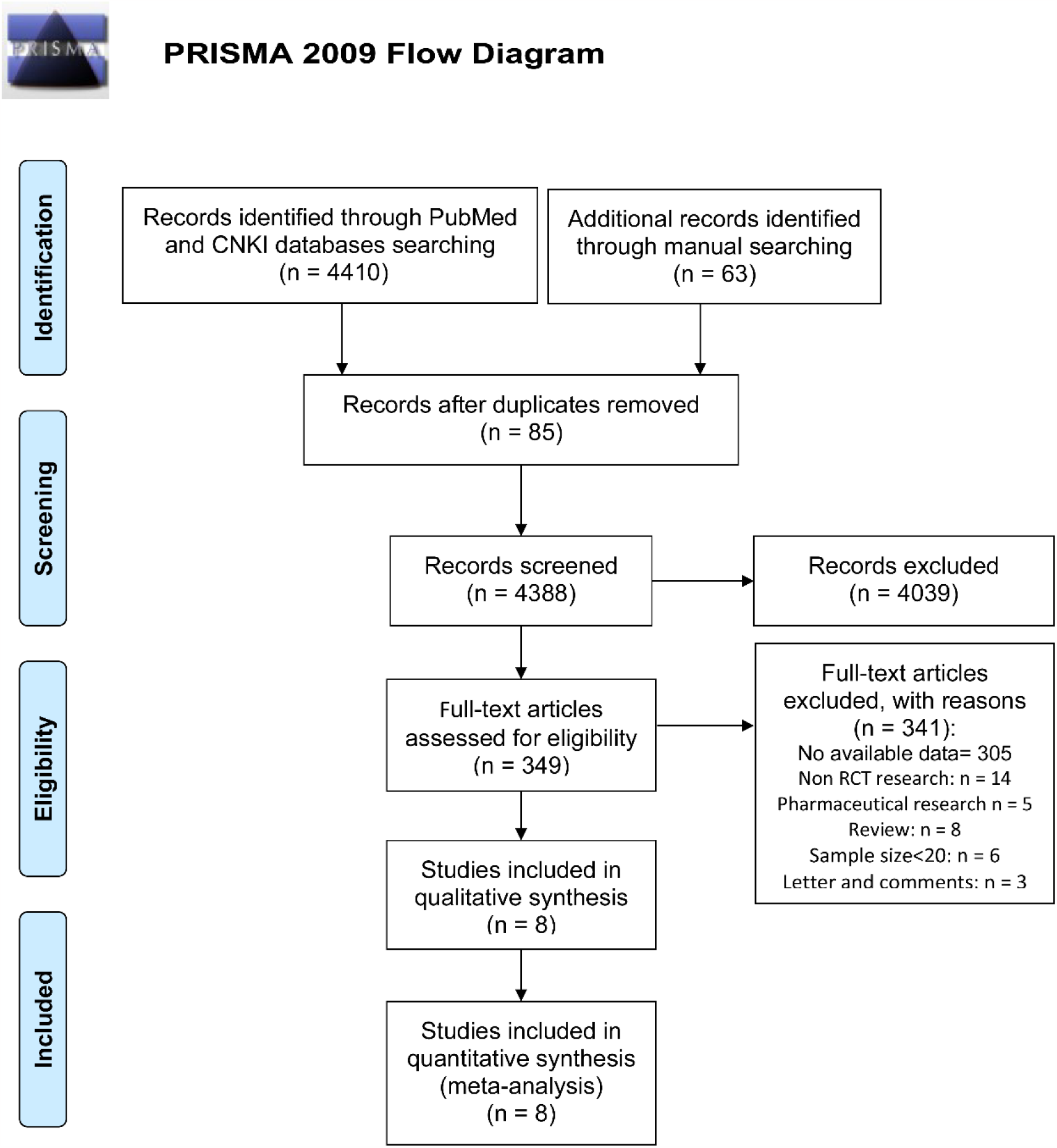
Study selection. RCTs, randomized clinical trials.

The meta-analysis showed that the use of PBL significantly improved problem-solving ability (SMD = 1.12, 95% CI = 0.25-1.99; **Figure 2A**) but had a high heterogeneity (I^2^ = 87%, *P*_heterogeneity_ < 0.01). PBL was also associated with better performance in self-directed learning (SMD = 1.40, 95% CI = 0.56-2.25; **Figure 2B**) and critical thinking (SMD = 1.55, 95% CI = 0.64-2.45; **Figure 2C**) but with high heterogeneity (both *P* < 0.01). However, there was no significant difference in the final exam score between the PBL group and the LBL control group (SMD = 0.23, 95% CI = -0.08-0.53; **Figure 2D**).

**Figure 2.**
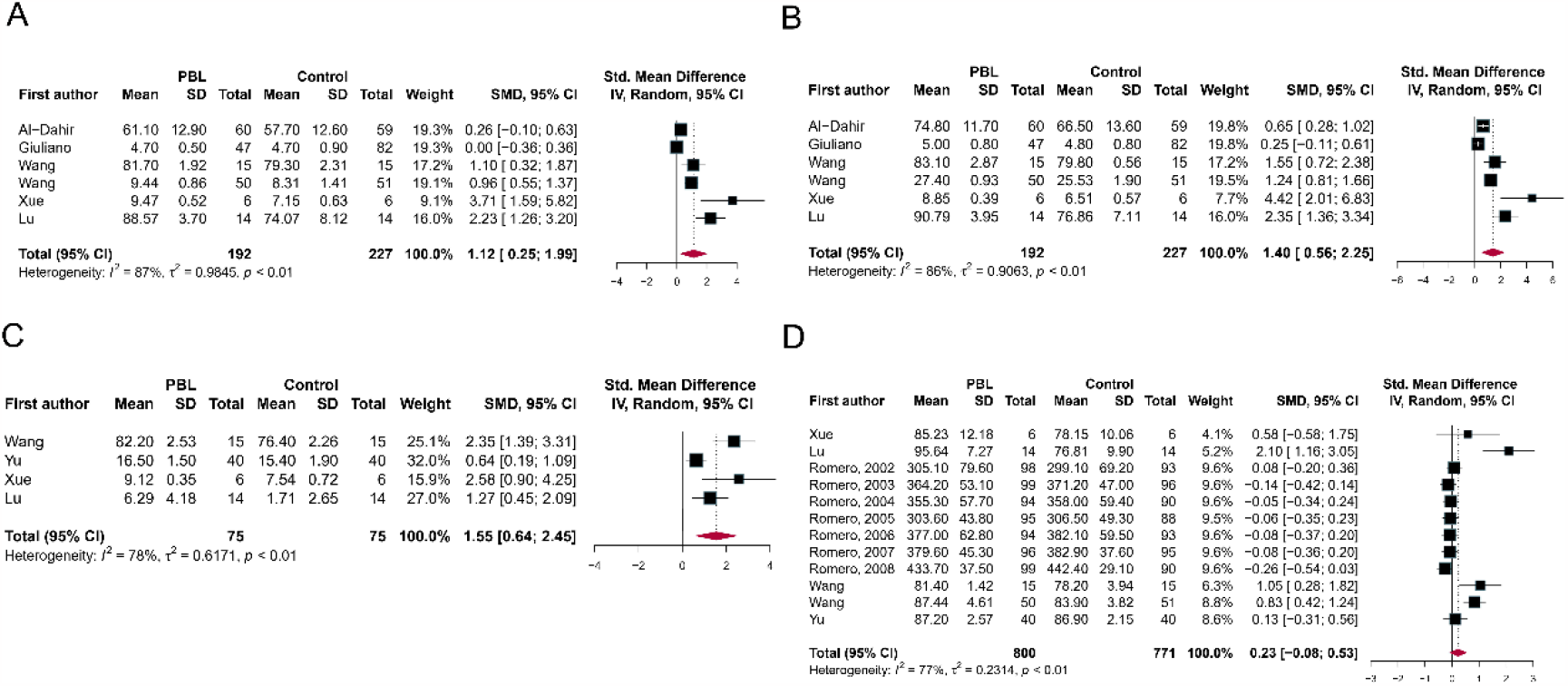
Meta-analysis for the effect of PBL compared with LBL group. (A) Problem-solving, (B) Self-directed learning, (C) Critical thinking, (D) Final exam score. PBL, problem-based learning; LBL, lecture-based learning.

A sensitivity analysis was conducted to assess the robustness of the meta-analysis results by removing each study in turn. The pooled results for problem-solving (**Supplementary Figure 1A**), self-directed learning (**Supplementary Figure 1B**), critical thinking (**Supplementary Figure 1C**), and final exam scores (**Supplementary Figure 1D**) were not materially changed by any of the removed studies. Furthermore, no significant asymmetry was observed in the funnel plot (**Supplementary Figure 2**). These findings suggest that the present meta-analysis is stable and reliable.

## DISCUSSION

Our study found that problem-based learning (PBL) significantly improves the performance of pharmacy students in problem-solving, self-directed learning, and critical thinking. The students’ feedback suggested that the pharmaceutics course based on PBL left them with broad knowledge that they were able to access long after the course ended. They also stated that the course encouraged the collaborative nature of the “real world,” self-directed learning orientation, and stimulated critical thinking. Subsequent meta-analysis confirmed our findings, indicating that PBL enhances students’ learning ability.

The steps of PBL include: (1) identifying the problem, (2) exploring prior knowledge, (3) generating potential theories, (4) identifying learning needs, (5) individually gathering research, (6) re-evaluating and applying new knowledge to the problem, and (7) reflecting on the learning process ^21^. PBL has been introduced into pharmacy education to prepare future pharmacists to meet the challenging requirements of the pharmacy profession^22, 23^. For our course in the China Pharmaceutical University, Authentication of Chinese Medicines, it is an applied discipline that aims to identify and study the varieties and quality of Chinese medicines, formulate standards for Chinese medicines, and find and expand new sources of medicines. Based on the inheritance of traditional Chinese medicine heritage and traditional identification and its application experience, this course applies modern scientific knowledge, techniques, and methods to research and focus on the theories and practices of the sources, properties, microscopic characteristics, physical and chemical identification, quality standards, and resources and sustainable utilization of Chinese medicines. The study of this course will lay the foundation for students to engage in authenticity identification, variety sorting, quality evaluation, development, and application of Chinese medicine after graduation, and ensure the safety and effectiveness of clinical medication. Because of the constantly evolving nature of the pharmacy profession, the skills acquired from PBL are necessary, as career opportunities have expanded to incorporate pharmacists as a pivotal partner for patient care In our cohort, we observed that PBL significantly increased the score of final exams.

However, this conclusion was not supported by the meta-analysis. One possible reason for this discrepancy is that our score included the normal performance score, which was higher in the PBL group due to the students’ enthusiastic speech and active participation in answering questions. Nonetheless, for problem-solving, self-directed learning, and critical thinking abilities, both our results and the meta-analysis demonstrated a significant improvement in the performance of pharmacy students who underwent PBL. These skills are more essential for future work than the final exam score.

In conclusion, our study provides evidence that PBL is an effective teaching method for pharmacy students, enhancing their problem-solving, self-directed learning, and critical thinking abilities. The impact of PBL on the career of a medicine student is profound and long-lasting.

## Declaration of interests

The authors have nothing to disclose.

## Notes

### Competing Interest Statement

The authors have declared no competing interest.

